# A pan-cancer atlas of transcriptional dependence on DNA methylation and copy number aberrations

**DOI:** 10.1101/2020.05.04.076901

**Authors:** Christian Fougner, Elen K. Höglander, Tonje G. Lien, Therese Sørlie, Silje Nord, Ole Christian Lingjærde

## Abstract

Cancer transcriptomes are shaped by genetic and epigenetic features, such as DNA methylation and copy number aberrations. Knowledge of the relationships between gene expression and such features is fundamental to understanding the basis of tumor phenotypes. Here, we present a pan-cancer atlas of transcriptional dependence on DNA methylation and copy number aberrations (PANORAMA). Our analyses suggest that copy number alterations are a central driver of inter-tumor heterogeneity, while the majority of expression-methylation associations found in cancer are a reflection of cell-of-origin and normal cell admixture. The atlas is made available through an online tool at https://pancancer.app.

## Main

The phenotype of a tumor is encoded in its transcriptome, which in turn is shaped by the tumor’s genome and epigenome. Changes in DNA methylation and copy number are frequently assumed to have a phenotypic effect. However, there is major variation, between genes and tumor types, in how strongly these features affect transcription, and consequently, their functional significance ^1,2,3^. The relationship between genetic factors and gene expression is essential to understanding transcriptomic regulation and heterogeneity in cancer, yet it is not well characterized at the whole-genome or pan-cancer level. Methods for interrogating these relationships are not yet adequately developed and large datasets are required. In this study, we develop tools for probing expression-methylation (E-M) and expression-copy number (E-C) associations, which we use to generate a pan-cancer atlas of transcriptional dependence on DNA methylation and copy number aberrations (Fig. 1a). The atlas is made available through a web application and is used to uncover distinct patterns of transcriptional regulation across genes and tumor types.

**Figure 1:**
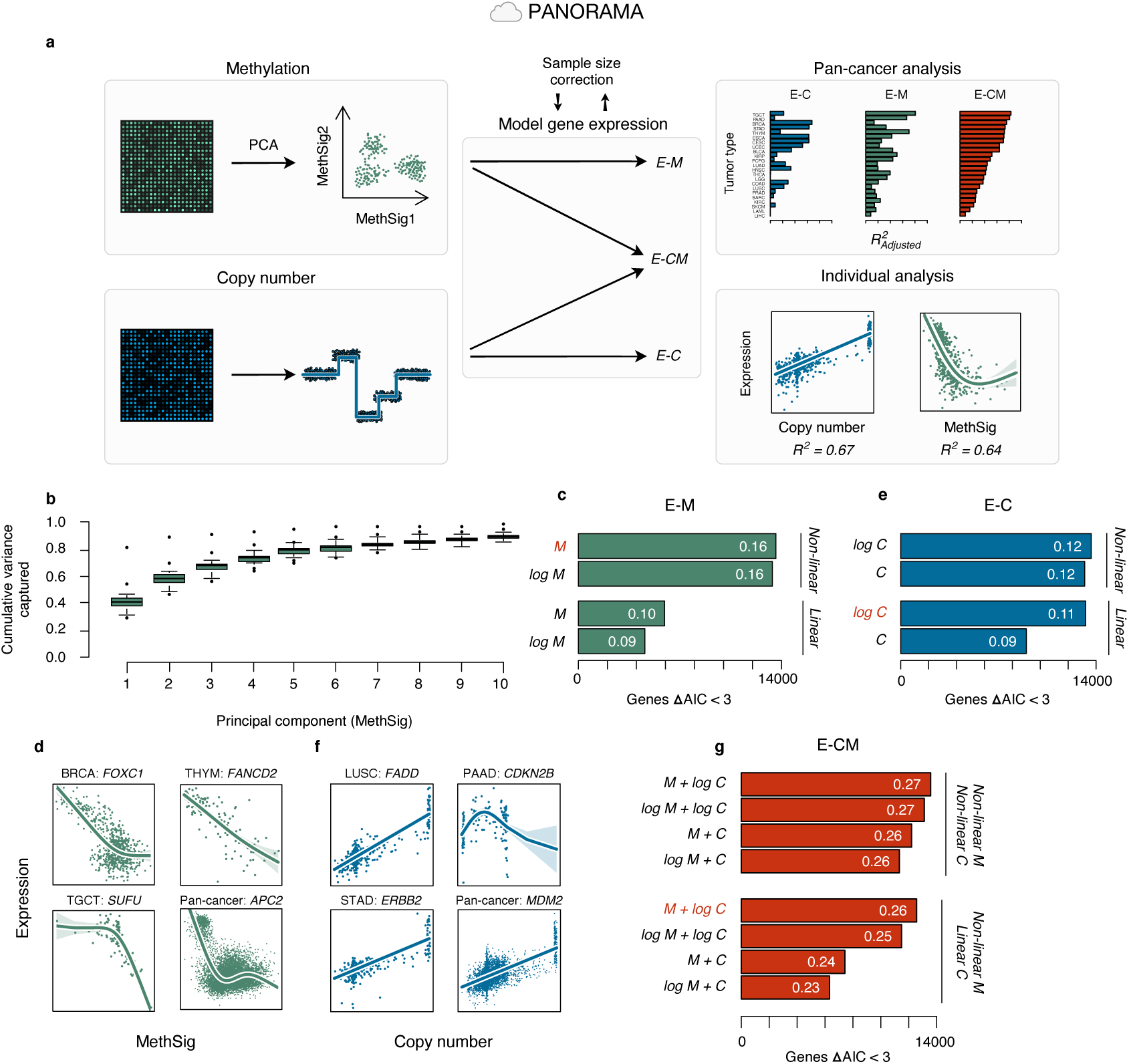
A framework for modeling the effect of differential DNA methylation and copy number on the cancer transcriptome. **a** Schematic of the analytical pipeline. PANORAMA is a pan-cancer atlas of expression-methylation and expression-copy number associations, integrated in an online tool for customized analyses. Analyses are based on The Cancer Genome Atlas pan-cancer dataset ^4^. **b** Median cumulative variance captured by principal components (MethSigs) 1-10 across 23 tumor types. Boxplot elements: center line = median, box limits = upper and lower quartiles, whiskers = 1.5 × interquartile range. **c - e** Gene expression modeled as a function of methylation. **c** Number of genes either modeled best using the given model (i.e. the model with the lowest AIC) or not modelled significantly worse than the best model (i.e. ΔAIC <3). Numbers inside the bars represent mean adjusted *R*^2^ for the model. **d** E-M association in selected genes, revealing diverse linear and non-linear dynamics. **e - f** Gene expression modeled as a function of copy number. **e** Number of genes with ΔAIC <3, and mean adjusted *R*^2^. **e** E-C association in selected genes, mainly revealing linear dynamics and occasional, potentially spurious, non-linear dynamics (e.g. *CDKN2B* in PAAD). Visualized with log-transformed copy number data. **g** Number of genes with ΔAIC <3, and mean adjusted *R*^2^, for gene expression modeled as a combined function of methylation and copy number. Linear methylation terms in E-CM models were also explored, but were excluded from the visualization. **All** Non-linear models were generated using splines with *k* = 4 basis functions in generalized additive models. Linear models generated using linear regression. **c, e, g** BRCA shown as a representative tumor type. Pan-cancer results shown in Supplementary Fig. 2. Models shown in red text were used onward.

In order to investigate the interplay between gene expression, methylation and copy number, all data levels first needed to be numerically represented in forms suitable for statistical modeling. The expression and copy number of a gene can be represented as scalars (i.e. a single numeric value; alternative RNA splicing and copy number breakpoints within coding regions are not considered here). In contrast, the methylation state of numerous CpGs may be relevant to the expression of a gene. To reduce the dimensionality of methylation data, and to achieve consistency across genes, we applied principal components analysis (PCA) to all CpGs within a window centered around each gene’s coding region. In the reported analyses, this window starts and ends 50,000 bases upstream and downstream of the respective genes coding region, thereby creating a broad representation of each gene’s *cis* methylation status. Across 23 tumor types in The Cancer Genome Atlas (TCGA)^4^, a median of 41% of all per-gene variation in methylation could be captured by the first principal component, and a median of 78% of variation could be captured with five principal components (Fig. 1b, Supplementary Fig. 1, Supplementary Data 1). Based on the variation captured, five principal components were selected as an appropriate trade-off between dimensionality reduction and information loss when modeling E-M associations. Methylation data transformed by PCA are onwards referred to as Methylation Signatures (MethSigs).

We next aimed to establish the dynamics of E-M and E-C associations in cancer. For this purpose, we considered both untransformed and log-transformed covariates, and linear and non-linear (non-parametric) ^5^ functional relationships. To identify the optimal model, we applied the Akaike Information Criterion (AIC) which quantifies the goodness-of-fit to the data while penalizing for high model complexity. Here, a model was considered significantly better than another if there was a difference in AIC of three or greater ^6^. Our analyses revealed a considerable improvement in modeling of E-M associations when allowing for non-linear relationships, but there was no improvement from log-transformation of MethSigs (Fig 1c, Supplementary Fig. 2a, Supplementary Data 2). Inspection of selected genes with strong E-M association showed varied, linear and non-linear relationships (Fig. 1d). In contrast, inclusion of non-linear terms in E-C models did in most cases not markedly improve the model fit, provided the copy number data were log-transformed (Fig. 1e, Supplementary Fig. 2b). Inspection of E-C associations in individual genes confirmed that relationships were mostly linear (Fig. 1f). A minority of E-C associations showed improved fit from inclusion of non-linear terms (e.g. *CDKN2B* in PAAD, Fig. 1f), however it is challenging to ascertain at genome-wide scale whether such non-intuitive relationships are spurious (i.e. overfitting) or representative of genuine biological features. Finally, we generated a combined model of gene expression as a function of copy number and methylation (E-CM). The combined model confirmed the necessity of non-linear methylation terms and that linear terms were adequate for copy number data if log-transformed (Fig. 1g, Supplementary Fig. 2c). These models illustrate a threshold effect whereby methylation correlates with transcription only within a relatively limited dynamic range. In contrast, there was no evidence of saturation effects in E-C associations at higher copy number levels (although copy number observations are here, to an extent, limited by SNP-array technology and data analysis methods ^7^).

We wished to generate a pan-cancer atlas of E-M/E-C associations allowing comparisons across tumor types. To handle the highly variable samples sizes across tumor types in TCGA ^4^, we randomly downsampled to 100 tumors per tumor type, modeled associations, and repeated this process 100 times. The median model statistics from these repeated runs were used as the estimate for the tumor type. Tumor types with less than 100 samples were removed from consideration (Supplementary Figure 3).

The pan-cancer atlas of transcriptional dependence on DNA methylation and copy number aberrations is made available through a web application at https://pancancer.app. PANORAMA provides, for each gene, an overview of transcriptional associations pan-cancer, while also enabling detailed analyses in individual tumor types or in a tissue-agnostic manner (Fig. 1a, 1d and 1f).

Analyses can be fully customized, including whether or not non-linearities are allowed when modeling, the number of MethSigs to use, and the size and location of the CpG window (thereby enabling *trans* analyses). PANORAMA generates publication-ready vector graphics, and provides raw and processed data for further analysis.

A survey of the pan-cancer atlas (Supplementary Data 3) revealed marked differences between tumor types in degree of transcriptional association to methylation and copy number. Genome-wide transcriptional association to copy number ranged from 2% in THCA to 14% in LUSC (Fig. 2a, Supplementary Data 2). There was a strong correlation between mean genomic instability index in a tumor type and mean E-C association (*R*^2^ = 0.64, *P* < 0.001, linear regression; Fig. 2b). Similarly, mean gene-centric copy number variance in a tumor type (an alternate measure for the aggregate burden of copy number aberration) was closely correlated to mean E-C association (*R*^2^ = 0.88, *P* < 0.001, linear regression; Supplementary Figure 4a). There was also a positive correlation between copy number variance and E-C association in individual genes (Supplementary Fig. 5). This correlation was, however, modest and genes with high copy number variance, but low E-C association abounded. These trends indicate that, at the whole-genome level, the burden of copy number aberration captures the extent to which a tumor transcriptome is copy number-driven reasonably well. At a single gene level though, there is considerable variation in transcriptional response to copy number aberration, illustrating the importance of analyzing copy number in the context of gene expression, e.g. with PANORAMA. Finally, there was significant correlation between mean gene expression variance in a tumor type and mean E-C association (*R*^2^ = 0.39, *P* = 0.002, linear regression; Supplementary Fig. 4b), and between mean copy number variance and mean gene expression variance (*R*^2^ = 0.49, *P* < 0.001, linear regression; Supplementary Fig. 4c). In sum, these correlations implicate copy number aberration as an important driver of transcriptional heterogeneity within tumor types.

**Figure 2:**
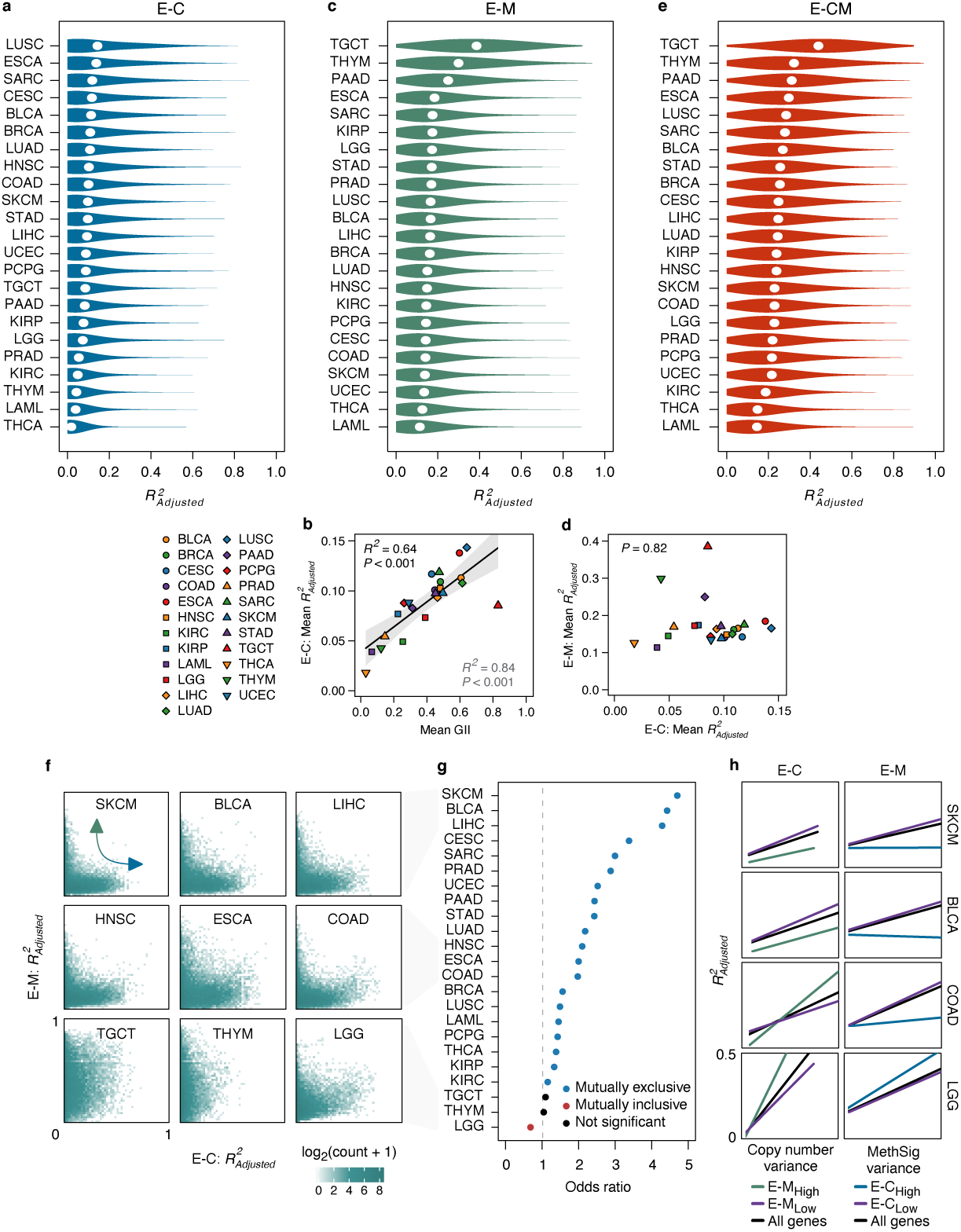
Pan-cancer patterns of transcriptomic association to DNA methylation and copy number. **a** Violin plots showing distribution of adjusted *R*^2^ for E-C associations across the genomes of 23 tumor types (ordered by mean adjusted *R*^2^, represented by white dots). **b** Correlation between mean genomic instability index (GII) in a tumor type and mean adjusted *R*^2^ for E-C associations (*R*^2^ = 0.64, *P* < 0.001, linear regression; *R*^2^ = 0.84, *P* < 0.001 if TGCT is treated as an outlier). Shaded dark grey area represents 95% confidence interval. **c** Violin plots showing distribution of adjusted *R*^2^ for E-M associations. **d** Correlation between mean adjusted *R*^2^ for E-C association and mean adjusted *R*^2^ for E-M association (*P* = 0.82, linear regression). **e** Violin plots showing distribution of adjusted *R*^2^ for E-CM associations. **F** 2D histograms (50 × 50 bins) showing adjusted *R*^2^ for E-C and E-M relationships in individual genes, in selected tumor types. In many tumor types, there appeared to be an anti-correlation between E-M and E-C association in individual genes. **g** Odds ratio for high E-C association and high E-M association in individual genes. Odds ratio was derived by separating genes into 2 × 2 contingency tables for genes with adjusted *R*^2^ above/below the 80^th^ percentile for E-M and E-C association. In most tumor types, there was a significant mutual exclusivity in whether genes showed high E-M or high E-C association (*P* < 0.05, Fisher’s exact test). Different percentile cut-offs are shown in Supplementary Figure 7. **h** Linear regresssion of adjusted *R*^2^ for E-C association correlated to copy number variance (left), and adjusted *R*^2^ for E-M association correlated to MethSig variance (right), in individual genes in selected tumor types. Linear regression was performed for all genes, and separately for genes stratified by E-M association above/below the 80^th^ percentile (E-M_high_/E-M_low_, left) and stratified by E-C association above/below the 80^th^ percentile (E-C_high_/E-C_low_, right). Full figures, including regression statistics, for all tumor types provided in Supplementary Figs. 5 and 6.

Genome-wide transcriptional association to methylation ranged from 11% in LAML to 39% in TGCT (Fig. 2c, Supplementary Data 2). TGCT, THYM and PAAD stood out as tumor types with exceptionally strong E-M associations, while the remaining tumor types showed relatively limited differences in mean E-M association. One explanation for these strong E-M associations might be extensive, systematic differences in methylation within certain tumor types, as exemplified by the divide between seminoma (globally hypomethylated) and non-seminoma in TGCT^8^. However, mean variance in MethSigs only correlated weakly with mean E-M association (*R*^2^ = 0.21, *P* =, linear regression; Supplementary Fig. 4d), and this correlation was entirely driven by one tumor type (TGCT). THYM and PAAD showed below-average MethSig variance. At a per-gene level, there were positive, but minor correlations between MethSig variance and E-M association (Supplementary Fig. 6). There was no association between mean gene expression variance in a tumor type and mean E-M association (*P* = 0.92, linear regression; Supplementary Fig. 4e). If aberrant methylation is a causal driver of gross transcriptional dysregulation within tumor types, one might expect mean E-M association to show some correlation with mean gene expression variance. There was, however, a positive correlation between mean MethSig variance and mean gene expression variance (*R*^2^ = 0.43, *P* < 0.001, linear regression; Supplementary Fig. 4f). Viewed together with aforementioned trends (Supplementary Figs. 4d and 4e), it is uncertain whether this is indicative of a causal relationship, or underlying confounding factors. Others have, for example, noted associations between methylation changes, structural variations ^9^, and copy number aberrations ^2,10^ (a trend also suggested here, Supplementary Fig. 4g). Taken together, these correlations raise questions about whether methylation drives genome-wide transcriptional dysregulation in cancer (as appears to be the case for copy number aberrations), fine-tunes transcriptional patterns at a smaller scale, or mostly plays a passive role in maintaining transcriptional states ^2,11^. Interestingly, GO-term analyses ^12,13,14^ revealed considerable enrichment of immune-related processes in the 20% most highly methylation-associated genes in most tumor types (Supplementary Data 4). The bulk of E-M associations in cancer may therefore essentially be a reflection of cell-of-origin and non-tumor cell infiltration, rather than a reflection of cancer-specific and etiologically relevant features. It is well-documented that aberrant methylation may have an oncogenic effect mediated by epigenetic silencing of certain genes (e.g. *MLH1, BRCA1*) and by effects related to genomic instability ^15,10^; aberrant methylation does, however, not seem to be a direct mechanism causing gross transcriptional dysregulation. The direct oncogenic effect of aberrant methylation might then be conceptually more analogous to the effects of mutations in individual genes than the genome-wide effects caused by copy number aberrations.

There was no correlation between mean E-M and mean E-C association across tumor types (*P* = 0.82, linear regression; Fig. 2d). Note that the relative strengths of E-M and E-C associations are not directly comparable as they are based on different models.

Using the combined model (E-CM), mean transcriptional association to methylation and copy number ranged from 15% in LAML to 44% in TGCT (Figure 2e, Supplementary Data 2). The distribution of E-CM associations showed similarity to the distribution of E-M associations, likely due to the overweight of methylation covariates relative to copy number covariates in the E-CM model.

When the strengths of E-M and E-C associations in individual genes were compared, a statistically significant anti-correlation between the two associations emerged in most tumor types (*P* < 0.05, Fisher’s exact test at an 80^th^ percentile cut-off; Fig. 2f, g, Supplementary Fig. 7). There were differences between tumor types, with, e.g. SKCM, BLCA and LIHC showing high degrees of mutual exclusivity between strong E-M and strong E-C association (cut-off above/below the 80^th^ percentile; onward referred to as E-M_High_/E-C_High_ and E-M_Low_/E-C_Low_). One exception was LGG, in which E-M_High_ and E-C_High_ genes tended to coincide (i.e. were mutually inclusive). Significant correlations between E-M and E-C association, in individual genes, were not observed in TGCT and THYM.

In light of these trends, we returned to the correlation between MethSig/copy number variance and strength of E-M/E-C association in individual genes, and stratified the analyses according to the strength of the opposite association (i.e. the correlation between copy number variance and strength of E-C association was determined separately for E-M_High_ and E-M_Low_ genes, and *vice versa*; Fig 2h, Supplementary Figs. 5 and 6). In general, higher copy number variance was correlated with stronger E-C association, irrespective of strength of E-M association (although E-M_Low_ genes tended to have slightly stronger E-C association than E-M_High_ genes, except in LGG; Supplementary Fig. 5). In contrast, higher MethSig variance was, in most tumor types, correlated with stronger E-M association in E-C_Low_ genes, but not in E-C_High_ genes (Supplementary Fig. 6). Several tumor types did, however, not show this trend, including LGG, THYM and PCPG. These trends indicate that the mutual exclusivity in E-M and E-C associations may partially be explained by copy number aberrations being able to override modulating effects of methylation on gene expression. This would be in agreement with findings from Sun *et al.* ^2^ in which they arrive at similar conclusions based on analyses of conditional independence in expression, methylation and copy number. The trend of mutual exclusivity should also be viewed in light of the earlier discussion, in which we questioned the role of E-M associations in driving genome wide transcriptomic variation within tumor types (Supplementary Fig. 4) and the observation that E-M_High_ genes are frequently immune-related (Supplementary Data 4). The mutual exclusivity might then be explained by E-C_High_ genes primarily being those selected for in a cancer-associated evolutionary process ^16^ (Supplementary Data 5), whereas E-M_High_ genes are the result of distinct processes (e.g. normal-cell infiltration). It is therefore interesting to note there is a paucity of immune-related GO-terms enriched in E-M_High_ genes in LGG (Supplementary Data 4), indicating unique processes molding E-M associations in that tumor type (perhaps associated with hypermethylator phenotypes caused by IDH-mutations ^17^).

In sum, we have here generated novel methods and tools for investigating the origins of transcriptomic dysregulation in cancer. These tools were applied to a dataset of over seven thousand tumors, across 23 types of cancer, yielding significant insights into the interplay between methylation, copy number and gene expression.

## Materials and Methods

### Cohorts

The TCGA pan-cancer cohort was used in this study (*n* = 8213) ^4^. Only tumors with data available for all three data levels (gene expression, copy number, and methylation as measured by Illumina HumanMethylation450 arrays) were included. The various tumor types had the following sample sizes: Acute Myeloid Leukemia (LAML; *n* = 111), Adrenocortical Carcinoma (ACC; *n* = 76), Bladder Urothelial Carcinoma (BLCA; *n* = 398), Brain Lower Grade Glioma (LGG; *n* = 511), Breast Invasive Carcinoma (BRCA; *n* = 750), Cervical Squamous Cell Carcinoma and Endocervical Adenocarcinoma (CESC; *n* = 293), Cholangiocarcinoma (CHOL; *n* = 36), Colon Adenocarcinoma (COAD; *n* = 266), Esophageal Carcinoma (ESCA; *n* = 162), Glioblastoma Multiforme (GBM; *n* = 54), Head and Neck Squamous Cell Carcinoma (HNSC; *n* = 492), Kidney Chromophobe Renal Cell Carcinoma (KICH; *n* = 61), Kidney Renal Clear Cell Carcinoma (KIRC; *n* = 239), Kidney Renal Papillary Cell Carcinoma (KIRP; *n* = 257), Liver Hepatocellular Carcinoma (LIHC; *n* = 356), Lung Adenocarcinoma (LUAD; *n* = 428), Lung Squamous Cell Carcinoma (LUSC; *n* = 350), Diffuse Large B-cell Lymphoma (DLBC; *n* = 47), Mesothelioma (MESO; *n* = 83), Pancreatic Adenocarcinoma (PAAD; *n* = 169), Pheochromocytoma and Paraganglioma (PCPG; *n* = 164), Prostate Adenocarcinoma (PRAD; *n* = 486), Rectum Adenocarcinoma (READ; *n* = 87), Sarcoma (SARC; *n* = 246), Skin Cutaneous Melanoma (SKCM; *n* = 462), Stomach Adenocarcinoma (STAD; *n* = 364), Testicular Germ Cell Tumors (TGCT; *n* = 138), Thymoma (THYM; *n* = 119), Thyroid Carcinoma (THCA; *n* = 471) Uterine Carcinosarcoma (UCS; *n* = 56), Uterine Corpus Endometrial Carcinoma (UCEC; *n* = 387) and Uveal Melanoma (UVM; *n* = 80). In described analyses, only the 23 tumor types with at least one hundred samples were included. All tumor types, irrespective of sample size, can be analyzed through the PANORAMA web application (with the exception of Ovarian Serous Cystadenocarcinoma, for which there were Illumina HumanMethylation450 array data available for only ten samples).

### Gene Expression

Gene expression data, pre-processed as previously described ^4^, were retrieved from the TCGA Pancan Atlas publication page (see Data availability). In brief, data were generated by RNA-sequencing, and RSEM data were upper quartile normalized and batch corrected. Data were transformed to log_2_(*ReadCount* + 1). The standard deviation of each gene (calculated across all tumor types) was visualized as a density plot, and the distribution of standard deviations was found to be bimodal. Genes with standard deviation less than 0.195 were filtered out.

### Copy number

Per-gene copy number data, pre-processed as previously described ^4^, were retrieved from the TCGA Pancan Atlas publication page (see Data availability). In brief, copy number profiles were generated using Affymetrix SNP 6.0 arrays, segmented using circular binary segmentation ^18^, and made gene-centric using Ziggurat Deconstruction in GISTIC2.0^7^. Log-transformed copy number data were analyzed and visualized in the form of log_2_(*CopyNumber/*2). Due to known non-linear saturation effects in array probes and limitations of analytical methods, a cap was set at a maximum copy number of ∼25 (log_2_(*CopyNumber/*2) = 3.657) ^7^. In order to break ties when modeling, jitter was added to the smallest and largest possible data points.

To calculate genomic instability index, the number of nucleotides in copy number aberrant segments was divided by the total number of nucleotides in all segments. Segments were defined as copy number aberrant if their nearest integer copy number state deviated from their ploidy (calculated using ABSOLUTE ^19^). In tumors for which an ABSOLUTE solution was not available, ploidy was set to two.

### Methylation

Methylation data from Illumina HumanMethylation450 arrays, pre-processed as previously described ^4^, were retrieved from the TCGA Pancan atlas publication page (see Data availability). Missing values were imputed for each tumor type separately using the 10 nearest neighbors ^20^. Principal Component Analysis (PCA) was performed for each gene using zero-centered *β*-values for all CpGs within a window starting 50 000 base pairs upstream from the start of the coding region to 50 000 base pairs downstream from the end of the coding region. This window size was selected as a reasonable compromise between sensitivity and specificity, while window size and location can be freely selected in the accompanying web application. PCA was performed separately for each tumor type. The first five principal components (MethSigs) were used for E-M and E-CM modeling. Genes with fewer than five associated CpG probes were excluded from analyses in order to ensure that all models had the same number of covariates.

### Models

To investigate the relation between gene expression and methylation and copy number, a range of different regression models of varying complexity were employed. For a given gene and cancer type, let *M*_1_, *…, M*_5_ denote MethSig values (log-transformed or not), *C* copy number (log-transformed or not), and *E* log-transformed gene expression. Linear regression was used to fit *E* as a function of *M*_1_, *…, M*_5_, as a function of *C*, and as a function of both *M*_1_, *…, M*_5_ and *C*. The linear terms for *M*_1_, *…, M*_5_ and/or *C* were subsequently replaced by smooth non-parametric terms to form additive models ^5^ that allowed for non-linear effects of *M*_1_, *…, M*_5_, or of *C*, or of both. Linear models were fitted using the function lm in R, while additive models were fitted using the function gam in the R package *mgcv* ^21^. *Each smooth term was modeled as a regression spline with k* = 4, corresponding to at most three effective degrees of freedom (EDF). The actual EDF per smooth term is determined by model selection using the restricted maximum likelihood (REML) method, which is implemented in the gam function. To facilitate comparison of results across cancer types with highly variable sample sizes, we employed a strategy of repeated downsampling within each cancer type. For each gene, cancer type and choice of model (as described above) we fitted the model to 100 randomly chosen samples and computed model statistics. This procedure was repeated 100 times and median values for model statistics were used in further analyses. If more than 80% of the randomly chosen samples in a run had identical expression values for a gene, no model was generated for that gene in the given run. Negative adjusted *R*^2^ values were set to zero. Only protein coding genes with available data from all three data levels were analyzed. The Akaike Information Criterion (AIC) was used to evaluate the fit of each model, and for two models with a difference in AIC <3, the model with the smallest AIC value is considered to be substantially better ^6^. GO term enrichment analyses for biological processes were carried out using PANTHER ^12,13,14^. Enrichment was tested for using Fisher’s exact test with FDR correction (terms with FDR *P* < 0.05 listed in Supplementary Data 4 and 5).

## Code availability

All code used in the described analyses is available at: https://github.com/clfougner/PANORAMA

Code for the PANORAMA web application is available at: https://github.com/clfougner/PANORAMA_app

## Data availability

Data from the TCGA pan-cancer cohort ^4^ were queried from: https://gdc.cancer.gov/about-data/publications/PanCan-CellOfOrigin

and:

https://gdc.cancer.gov/about-data/publications/PanCan-Clinical-2018

The following files were used in the study:

Gene expression:

~~~
EBPlusPlusAdjustPANCAN_IlluminaHiSeq_RNASeqV2.geneExp.tsv
~~~

Methylation:

~~~
jhu-usc.edu_PANCAN_HumanMethylation450.betaValue_whitelisted.tsv
~~~

Per-gene copy number data:

~~~
ISAR_GISTIC.all_data_by_genes.txt
~~~

Copy number segments:

~~~
ISAR_corrected.PANCAN_Genome_Wide_SNP_6_whitelisted.seg
~~~

Ploidy:

~~~
TCGA_mastercalls.abs_tables_JSedit.fixed.txt
~~~

Clinical data:

~~~
TCGA-CDR-SupplementalTableS1.xlsx
~~~

Data from selected models are available in Supplementary Data 3. Full data from all generated models are available at: https://github.com/clfougner/PANORAMA/Output/AllModels.zip

## Acknowledgements

C.F. is supported by grants from the Norwegian Research Council (163027) and South-Eastern Norway Regional Health Authority (2012056) to T.S. S.N. was a researcher on a career grant from the South-Eastern Norway Regional Health Authority (Grant number 2014061), and E.K.H. was a postdoctoral fellow on the same grant. The authors would like to acknowledge the valuable scientific discussion and input from Anne-Lise Børresen-Dale and Katherine Hoadley.

## Author contributions

E.K.H, S.N. and O.C.L. conceived of and initiated the study. C.F., E.K.H, T.G.L., S.N. and O.C.L. designed the study. C.F., E.K.H. and S.N. carried out analyses. C.F., E.K.H, T.G.L., T.S., S.N. and O.C.L. interpreted results. C.F. made the web application. T.S., S.N. and O.C.L. supervised the study. T.S. and S.N. acquired funding. C.F. wrote the initial manuscript draft. C.F., E.K.H, T.G.L., T.S., S.N. and O.C.L. reviewed and edited the manuscript.

## Competing interests

E.K.H. is employed by Roche Norge AS since 01.10.2017. Roche Norge AS is a subsidiary of F. Hoffmann-La Roche Ltd. The research was conducted without any involvement by Hoffmann-La Roche and any personal views of E.K.H. should not be understood or quoted as being made on behalf of or reflecting the position of Hoffmann-La Roche. The remaining authors declare no competing interests.

## Supplementary Information

**Supplementary Figure 1:**
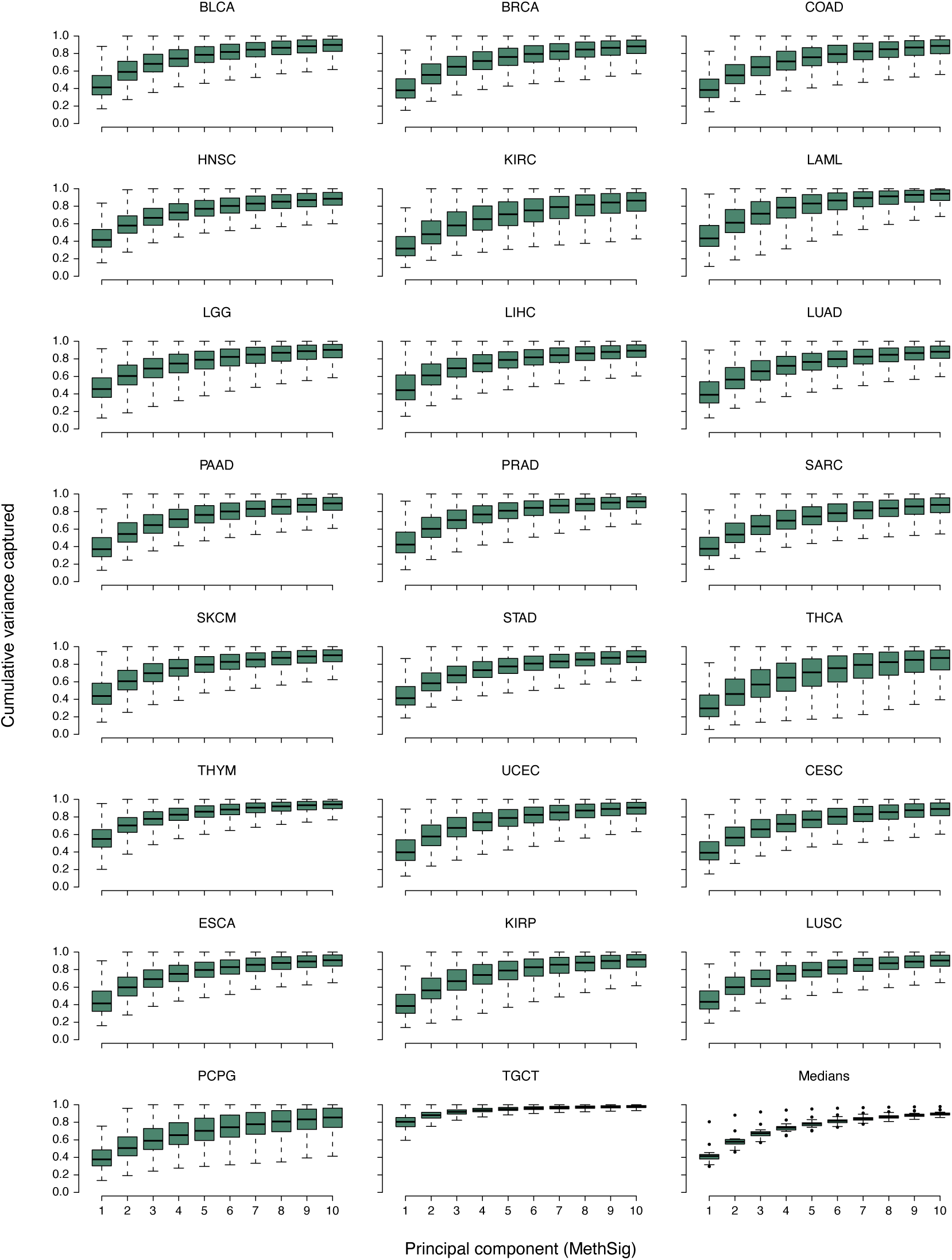
Dimensionality reduction of gene-centric methylation data using Principal Component Analysis (PCA). Boxplots of cumulative variance captured by PCA in gene-centric methylation data. PCA was performed on *β*-values for all CpGs within a window starting 50 kilobases upstream of the start of the coding region of a gene to 50 kilobases downstream of the end of the coding region. PCA was performed individually for each tumor type. Data points in the bottom right panel represent the median variances captured in the 23 tumor types in the TCGA pan-cancer dataset with at least 100 tumors (also shown in Fig. 1b). Boxplot elements: center line = median, box limits = upper and lower quartiles, whiskers = 1.5 × interquartile range; outliers only shown for the bottom right panel.

**Supplementary Figure 2:**
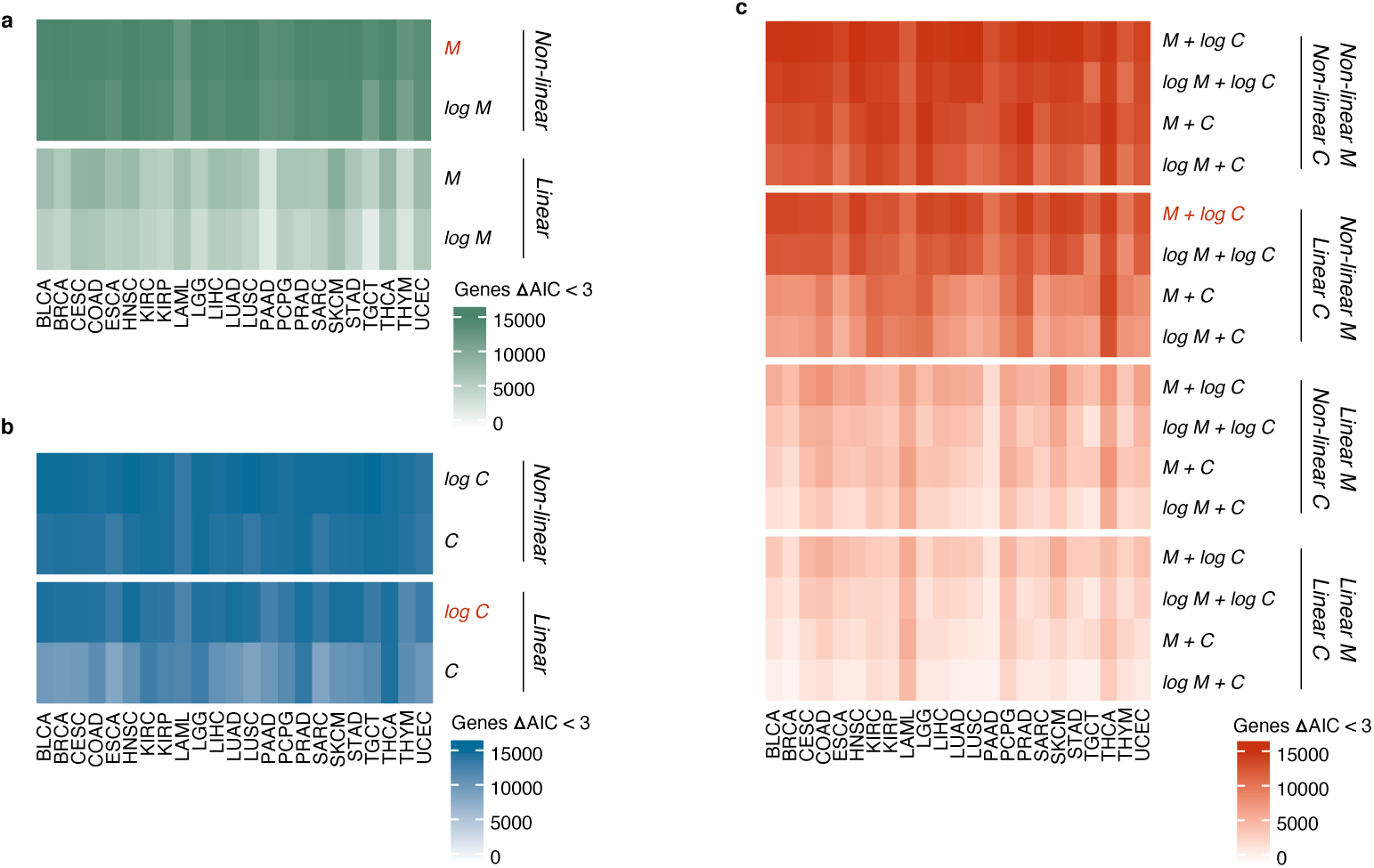
Modeling gene expression as a function of methylation and copy number. **a** Heatmap showing the number of genes with ΔAIC <3 for each E-M model. Gene expression modeled as a function of methylation showed marked improvement when non-linear spline terms were used, but no improvement was seen from log-transformation of MethSigs. **b** Heatmap showing the number of genes with ΔAIC <3 for each E-C model. Gene expression modeled as a function of copy number only showed minor, and potentially spurious, improvement from the addition of non-linear spline terms, provided copy number data were log-transformed. **c** Heatmap showing the number of genes with ΔAIC <3 for each E-CM model. The combined model confirmed that methylation data required non-linear spline terms for optimal modeling and that copy number was adequately described by linear terms if log-transformed.

**Supplementary Figure 3:**
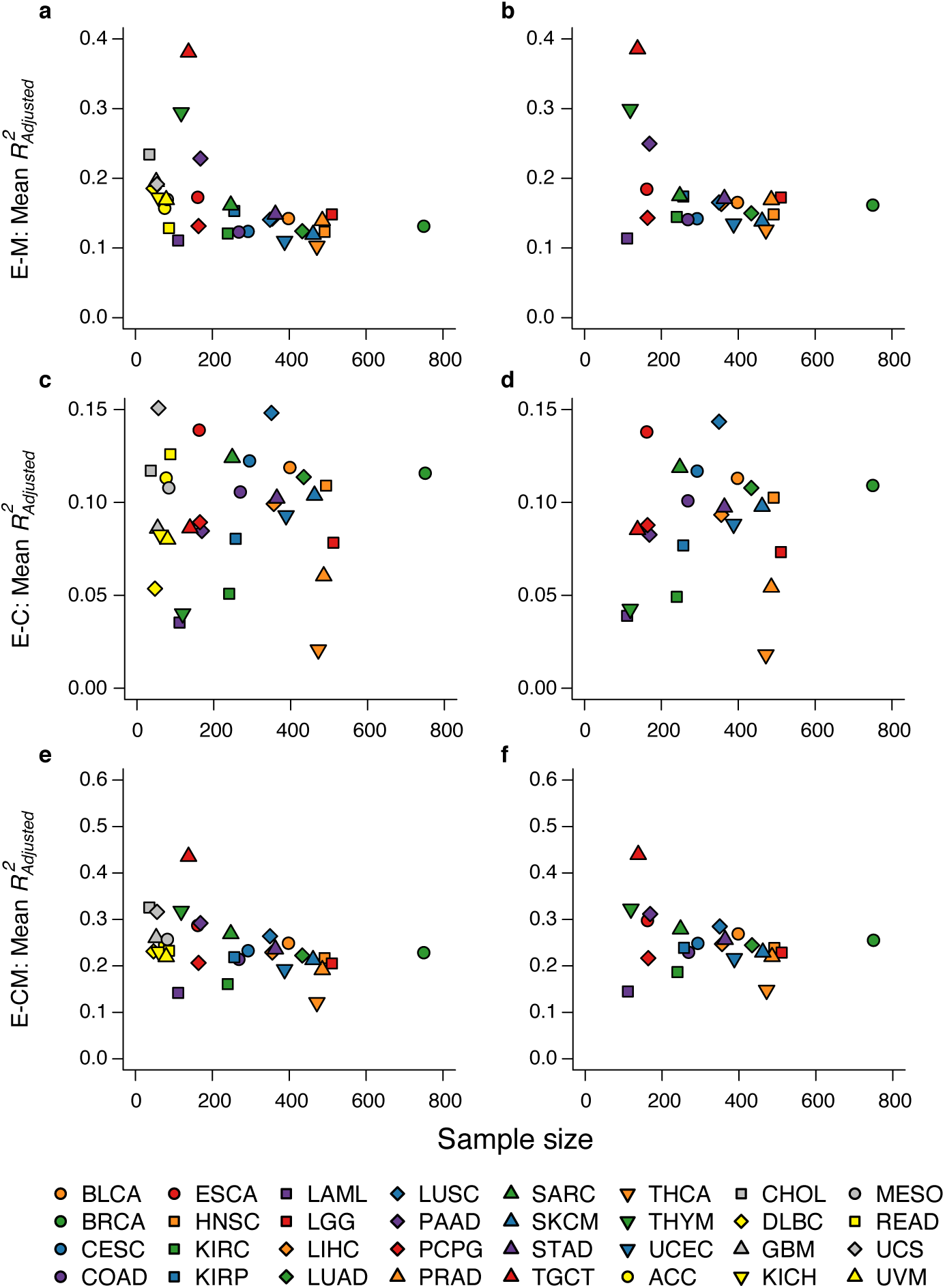
Mitigating sample size as a confounding factor. **a - b** Mean adjusted *R*^2^ for expression-methylation (E-M) associations before (**a**) and after (**b**) correction for samples size. **c - d** Mean adjusted *R*^2^ for expression-copy number (E-C) associations before (**c**) and after (**d**) correction for samples size. **e - f** Mean adjusted *R*^2^ for combined models (E-CM) before (**e**) and after (**f**) correction for samples size. **All** Sample size was corrected for by downsampling tumor types to one hundred samples, modeling transcriptional associations, then repeating downsampling and modeling one hundred times. The median model statistics from repetitions were used for further analyses. Sample size correction necessitated the exclusion of tumor types with fewer than one hundred samples.

**Supplementary Figure 4:**
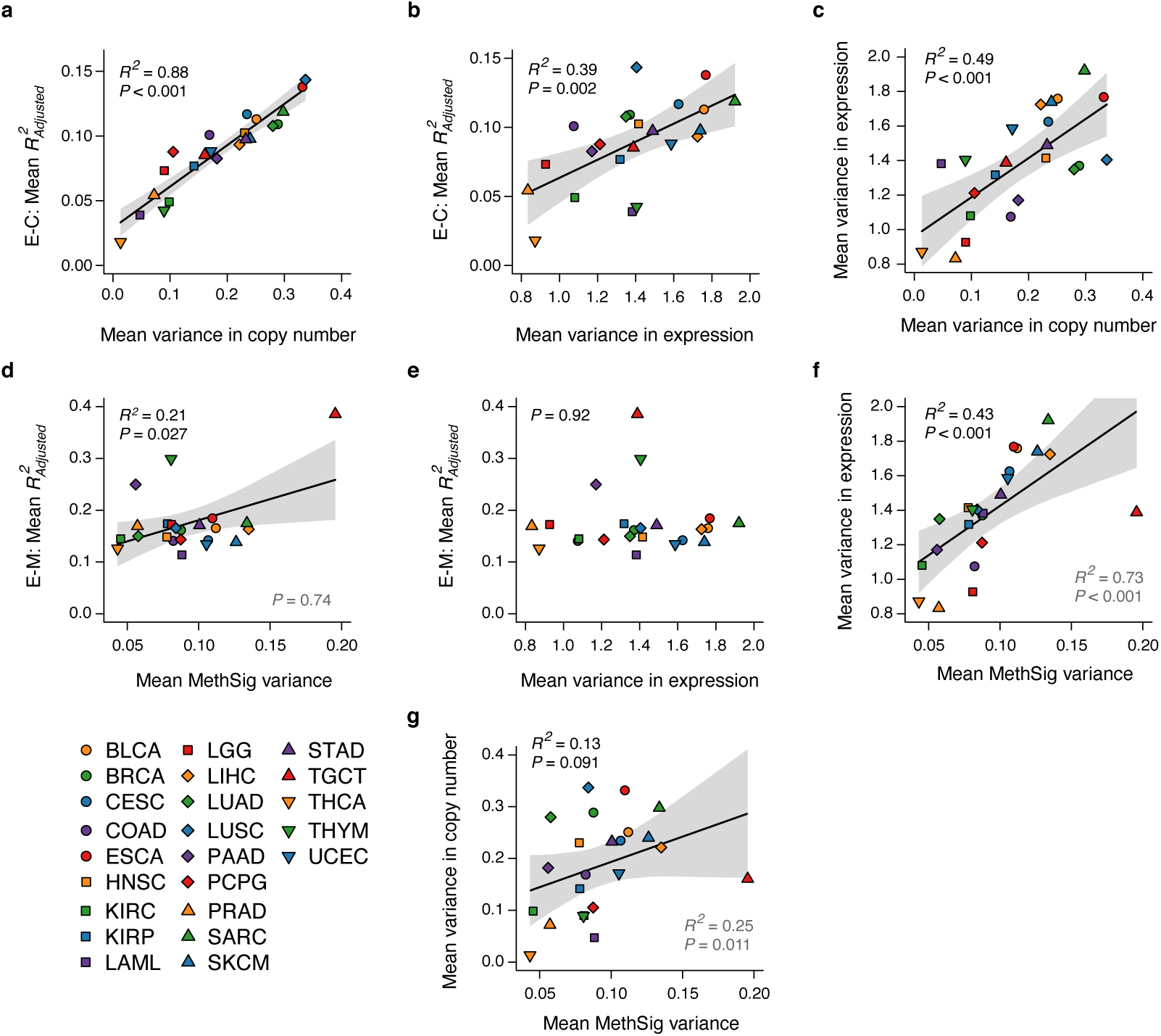
Importance of genome-wide variance in expression, copy number and methylation. **a** Correlation between mean variance in log-transformed, gene-centric copy number and mean adjusted *R*^2^ for E-C associations (*R*^2^ = 0.88, *P* < 0.001, linear regression). **b** Correlation between mean variance in log-transformed gene expression and mean adjusted *R*^2^ for E-C associations (*R*^2^ = 0.39, *P* = 0.002, linear regression). **c** Correlation between mean variance in log-transformed gene-centric copy number and mean variance in log-transformed gene expression (*R*^2^ = 0.49, *P* < 0.001, linear regression). **d** Correlation between mean MethSig variance (mean of MethSigs 1-5) and mean adjusted *R*^2^ for E-M associations. There was a positive correlation between the degree of variance in methylation in a tumor type and mean E-M association (*R*^2^ = 0.21, *P* = 0.027, linear regression), however this correlation is non-existent if TGCT is treated as an outlier (*P* = 0.74, linear regression, grey text). **e** Correlation between mean variance in log-scaled gene expression and mean adjusted *R*^2^ for E-M relationships (*P* = 0.92, linear regression). **f** Correlation between mean MethSig variance and mean variance in log-transformed gene expression. There was a positive correlation between the degree of variance in methylation and the degree of variance in gene expression (*R*^2^ = 0.43, *P* < 0.001, linear regression). This correlation is strengthened if TGCT is treated as an outlier (*R*^2^ = 0.73, *P* < 0.001, linear regression, grey text). **g** Correlation between mean MethSig variance and mean variance in log-transformed, gene-centric copy number. There was a positive, but non-significant correlation between the degree of variance in methylation and the degree of variance in copy number (*R*^2^ = 0.13, *P* = 0.091, linear regression, grey text). This correlation is significant if TGCT is treated as an outlier (*R*^2^ = 0.25, *P* = 0.011, linear regression, grey text). **All** Shaded, dark grey areas represent 95% confidence intervals.

**Supplementary Figure 5:**
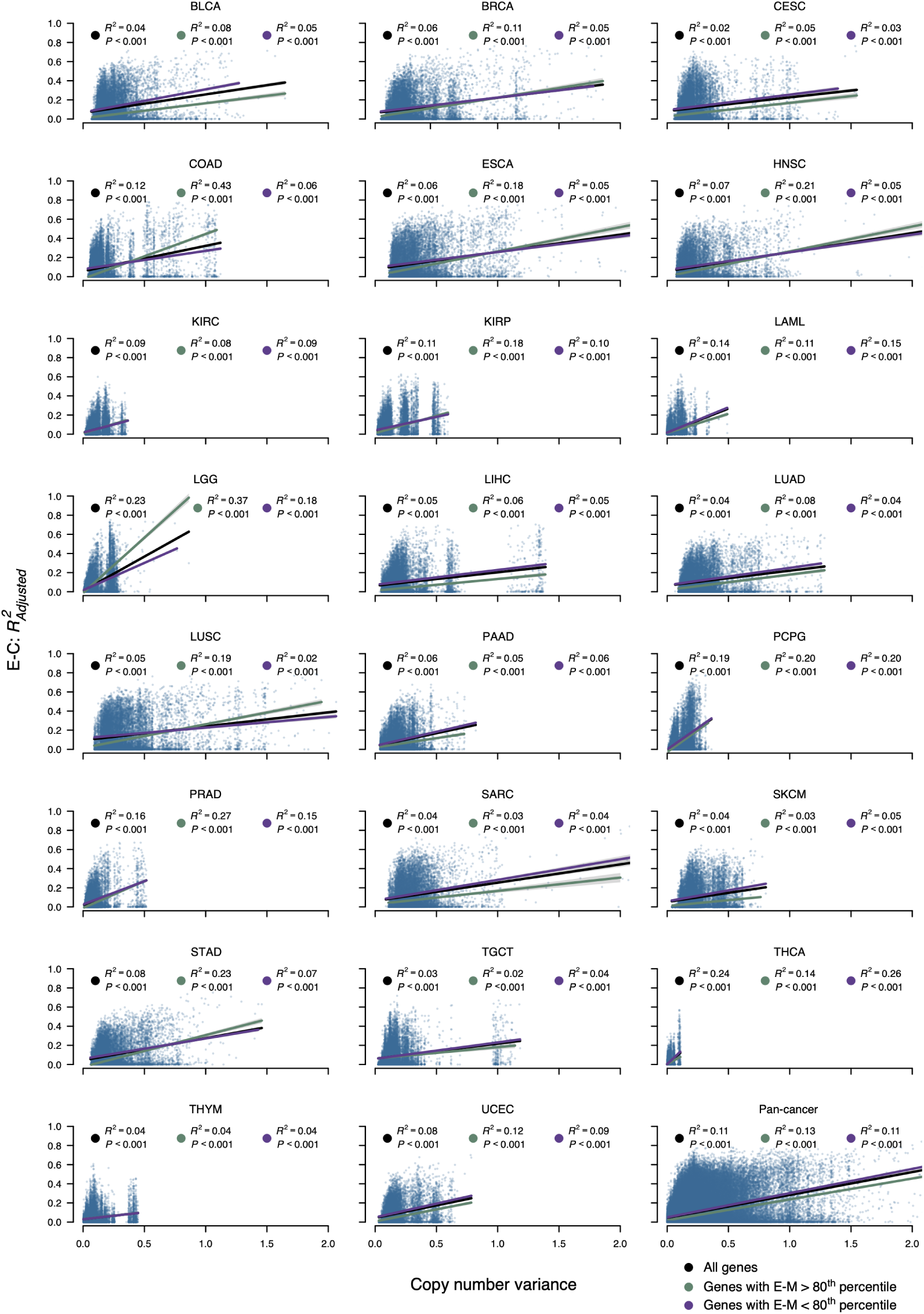
Importance of per-gene variance in copy number. Variance in copy number data plotted against E-C association at a per-gene level in 23 tumor types and pan-cancer. Linear regression was performed for all genes (black), and stratified by whether genes had E-M association above/below the 80^th^ percentile (green and purple, respectively). The pan-cancer plot represents the aggregate of the other 23 panels, and displays E-C association and copy number variance derived individually in each tumor type. The 80^th^ percentiles of E-M association used in the pan-cancer plot are identified individually for each tumor type.

**Supplementary Figure 6:**
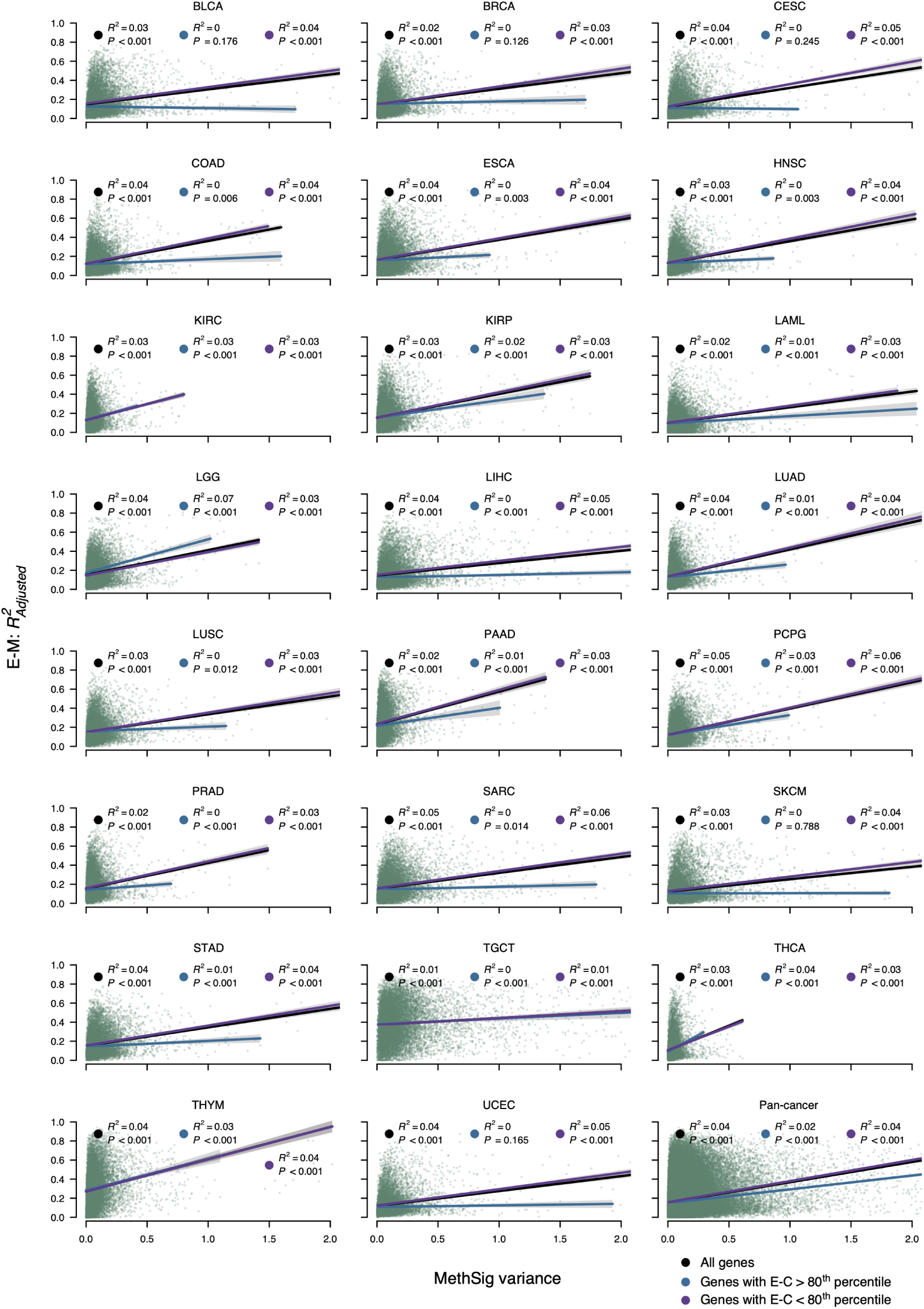
Importance of per-gene variance in MethSigs. Variance in MethSigs (mean of MethSigs 1-5) plotted against E-M association at a per-gene level in 23 tumor types and pan-cancer. Linear regression was performed for all genes (black), and stratified by whether genes had E-C association above/below the 80^th^ percentile (blue and purple, respectively). The pan-cancer plot represents the aggregate of the other 23 panels, and displays E-M association and MethSig variance derived individually in each tumor type. The 80^th^ percentiles of E-C association used in the pan-cancer plot are identified individually for each tumor type.

**Supplementary Figure 7:**
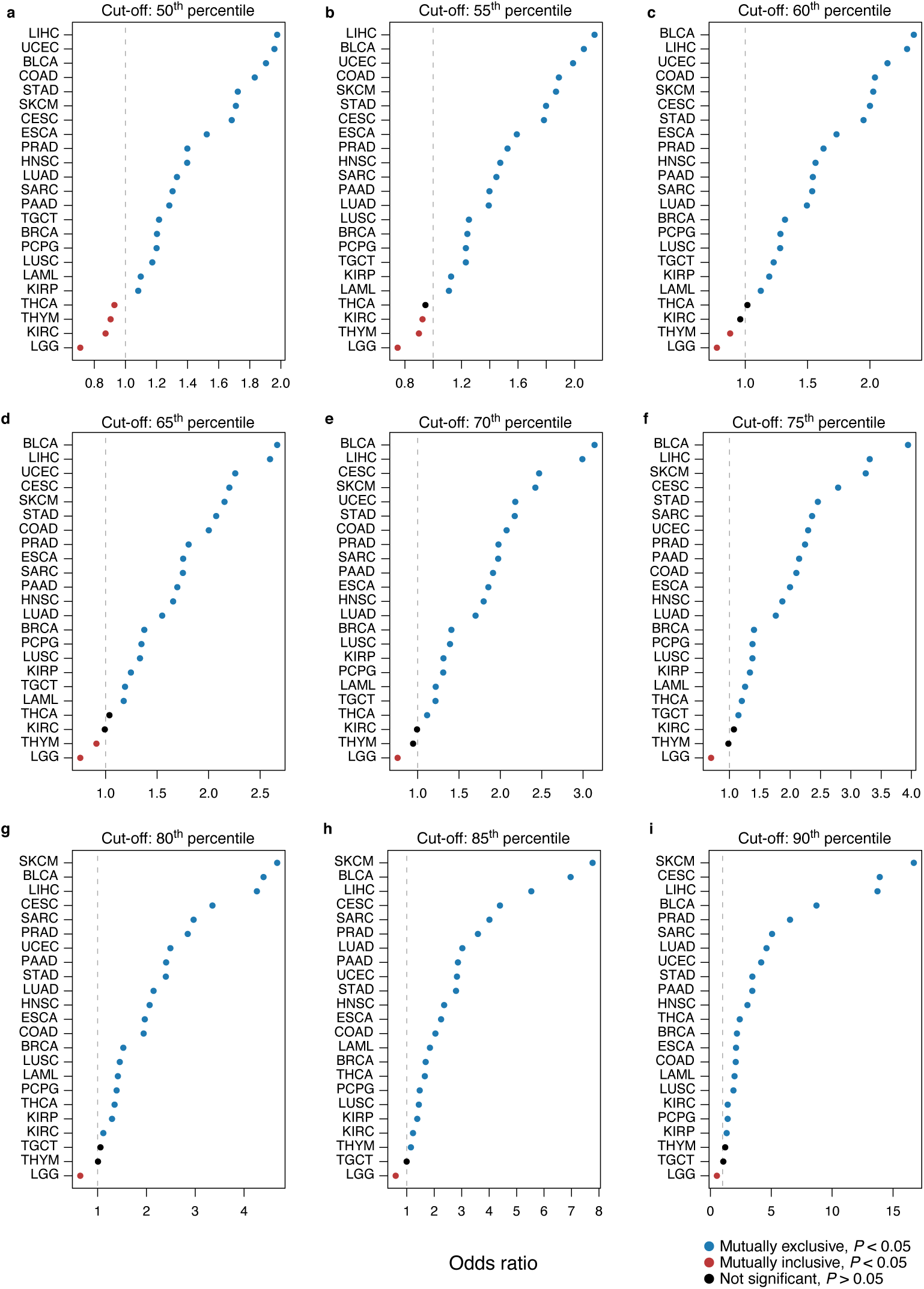
Mutual exclusivity in E-M and E-C association. Odds ratio for high level of E-M versus E-C association, defined at 50^th^ (**a**), 55^th^ (**b**), 60^th^ (**c**), 65^th^ (**d**), 70^th^ (**e**), 75^th^ (**f**), 80^th^ (**g**), 85^th^ (**h**) and 90^th^ (**i**) percentile cut-offs. **All** Odds ratio identified using 2 × 2 contingency tables of genes with adjusted *R*^2^ above/below the cut-off for E-M/E-C association. Significance derived using Fisher’s exact test. In most tumor types, genes with high level of one form of transcriptional association (E-M/E-C) showed significantly lower likelihood of high level of the other form of transcriptional association (E-C/E-M), i.e. the two associations were mutually exclusive. LGG was one exception in which E-M and E-C associations were mutually inclusive.

**Supplementary Data 1**: Median cumulative variance captured by principal components (Meth-Sigs) 1-10 in 23 tumor types (.xlsx).

**Supplementary Data 2**: Number of genes with ΔAIC <3, and mean adjusted *R*^2^, for tested models across 23 tumor types (.xlsx).

**Supplementary Data 3**: Genome-wide, pan-cancer model statistics for the 23 tumor types included in the pan-cancer atlas. (.xlsx).

**Supplementary Data 4**: Enriched GO-terms in the 20% most highly methylation-associated genes (E-M_High_) for each tumor type. (.xlsx).

**Supplementary Data 5**: Enriched GO-terms in the 20% most highly copy number-associated genes (E-C_High_) for each tumor type. (.xlsx).

